# Signatures of selection in a recent invasion reveals adaptive divergence in a highly vagile invasive species

**DOI:** 10.1101/643569

**Authors:** Adam P. A. Cardilini, Katarina C. Stuart, Phillip Cassey, Mark F. Richardson, William Sherwin, Lee A. Rollins, Craig D.H. Sherman

**Author notes:** Joint first Authors (Adam P. A. Cardilini, Katarina C. Stuart). Joint last authors (Lee A. Rollins, Craig D.H. Sherman). Corresponding Authors: Craig Sherman; Lee Ann Rollins.

## Abstract

A detailed understanding of population genetics in invasive populations helps us to identify drivers of successful introductions. Here, we investigate putative signals of selection in Australian populations of invasive common starlings, *Sturnus vulgaris*, and seek to understand how these have been influenced by introduction history. We use reduced representation sequencing to determine population structure, and identity Single Nucleotide Polymorphisms (SNPs) that are putatively under selection. We found that since their introduction into Australia, starling populations have become genetically differentiated despite the potential for high levels of dispersal, and that selection has facilitated their adaptation to the wide range of environmental conditions across their geographic range. Isolation by distance appears to have played a strong role in determining genetic substructure across the starling’s Australian range. Analyses of candidate SNPs that are putatively under selection indicate that aridity, precipitation, and temperature may be important factors driving adaptive variation across the starling’s invasive range in Australia. However, we also note that the historic introduction regime may leave footprints on sites flagged as being under adaptive selection, and encourage critical interpretation of selection analyses.

## Introduction

Understanding the proximate molecular mechanisms of natural selection is a central goal in evolutionary biology. Invasive species have long presented an opportunity to elucidate evolutionary mechanisms underlying selection (Baker and Stebbins 1965). Rapid adaptation is commonplace in invasive populations (Lee 2002; Rollins, et al. 2013) and is proposed to be important to their long-term success (Dlugosch and Parker 2008). The short evolutionary timescale over which this adaptation must take place provides opportunity to characterise the ecological and evolutionary processes that contribute to successful establishment and spread (Lee 2002). Understanding these proximate mechanisms enables us to predict how evolutionary adaptation will influence range distribution, or how populations (both native and invasive) may respond to environmental change (Bay, et al. 2018).

Investigating the genetics underlying evolutionary change in invasive populations gives us the opportunity to understand how a range of demographic and adaptive processes shape populations (e.g. bottlenecks, population expansions and adaptation) (reviewed in Sherman, et al. 2016). This information helps us understand invasion dynamics (Tepolt, et al. 2009) and may illuminate the adaptive response of species to a novel or changing environment (Riquet, et al. 2013). Using these data, we can identify signatures of molecular evolution in alien species (Ashburner, et al. 2000; Franks and Munshi-South 2014) by identifying loci putatively under selection when considering population genetic statistics such as F_ST_, or loci that are significantly associated with environmental variables (Foll and Gaggiotti 2008; Gunther and Coop 2013; Loh, et al. 2015).

Human introduced populations often have discrete (but potentially numerous and repetitious) geographic introductory sites, which will have variable numbers of founding individuals from potentially different source populations (Dlugosch and Parker 2008). What may then result is an amalgamation of subpopulations that, depending on the age of the populations and degree of connectivity, have the potential to be genetically differentiated geographically due to legacy genotypes (genotypes of founding individuals). Further, bottlenecks encourage stochastic evolution through processes such as genetic drift. During rapid range expansion, allelic surfing can create signatures at neutral loci that resemble selection, particularly at range edges (Klopfstein, et al. 2006; Lotterhos and Whitlock 2015; Hoban, et al. 2016; reviewed in Sherman et al. 2016). Until these discrete introductions make contact, it is possible that these processes are occurring concurrently and independently at different introduction sites and may leave genetic footprints. It is important to understand to what extent historic invasion regime influences patterns of adaptive genetic variation in invasive populations, particularly those with multiple and spatially separate introductions.

The common starling (*Sturnus vulgaris*) invasion in Australia represents an excellent system in which to assess the interaction between adaptive selection and historic introduction regime in shaping population structure. Starlings are a highly successful invasive species that has established populations around the world (Higgins, et al. 2006). Starlings were introduced into Australia from England during the 1850’s onwards at several locations, including Melbourne (Victoria, VIC), Brisbane (Queensland, QLD), Adelaide (South Australia, SA) and Hobart (Tasmania, TAS) (Fig. 1a), with the total individuals introduced numbering in the hundreds (Jenkins 1977; Higgins, et al. 2006). They quickly spread from their introduction sites on the south-eastern coastline to occupy a large range in south-eastern Australia (Long 1981, Rollins, et al. 2009; Rollins, et al. 2011), across habitats with significant environmental gradients (Woolnough, et al. 2005). They have now persisted in Australia for approximately 60 generations since their introduction (Higgins, et al. 2006; Rollins, et al. 2016).

**Fig. 1:**
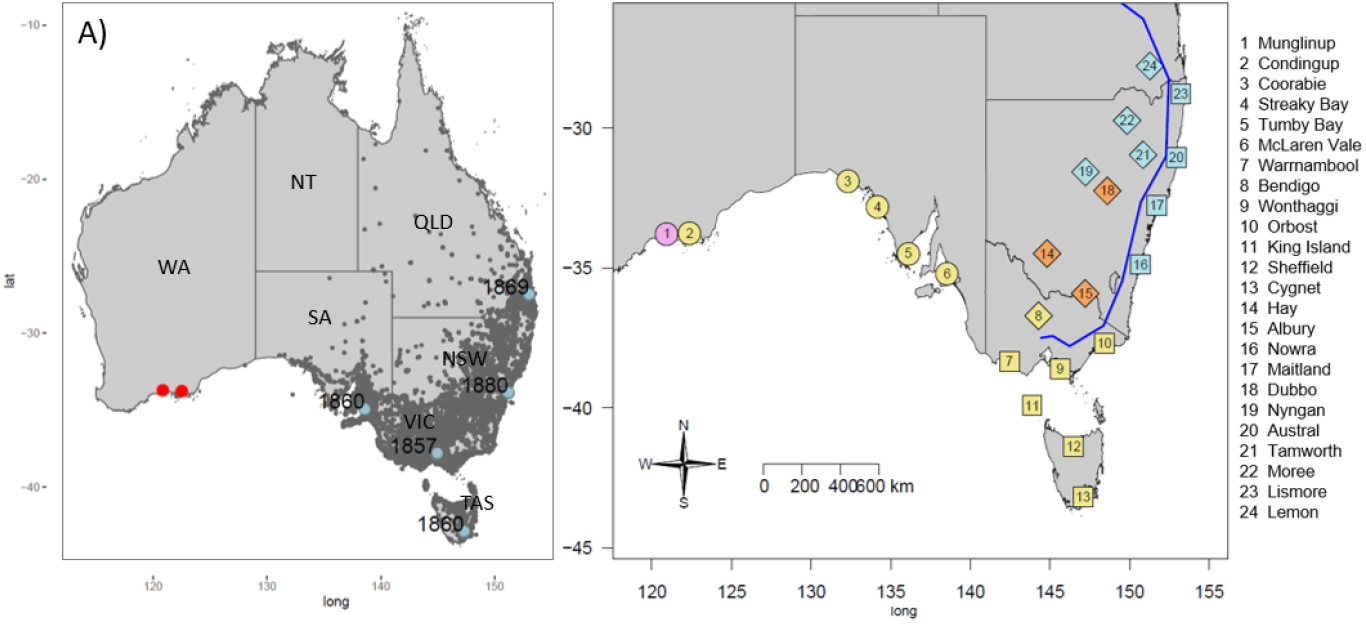
A) Australian starling distribution map according to eBird data (accessed Nov 2018), with introduction sites marked and first major introduction date listed adjacently, and current known range edge marked in red. B) Map of southeastern Australia. Points denote starling collection localities. Point shape indicates the environmental cluster that the point belonged to, as identified by a PCA analysis of environmental variables; circle = arid, diamond = semi-arid, square = non-arid. Point colour indicates the genetic cluster that the population belonged to as identified by population Bayesian fastSTRUCTURE analysis; pink = Munglinup cluster, yellow = southern Australia cluster, orange = southwest New South Wales cluster, and blue = northern Australia cluster. The blue line represents The Great Dividing Range (continental divide). Sample site number key corresponds to that listed in Table 1.

Starlings are predominantly found in agricultural landscapes relying on open fields to forage (Feare 1985; Whitehead, et al. 1995). The starling native range covers a wide range of environments spanning the Palearctic, including temperate forests and grasslands, as well as both hotter and colder climates at their native range edges (Higgins, et al. 2006). Starlings are a highly dispersive species, able to disperse upwards of 1000km from their natal site (Waterman, et al. 2008). Nevertheless, across their Australian range, starlings have two possible major barriers to movement. The hot, arid areas of Australia present a challenge to the species; central Australia is unlikely to be colonised by starlings and the large desert expanse of the Nullarbor Plain (between SA and WA) is known to restrict the species’ movement (Woolnough, et al. 2005). Second, large contiguous areas of forest and elevated mountain ranges along the Eastern coast of the continent are likely to act as a barrier to starling movement, restricting gene flow (Fig. 1b). High elevation is suspected to play an important role in shaping starling demography within starling invasive populations (Ross 1983), because the species has an upper elevation limit of 2000m (Higgins, et al. 2006). These biogeographical barriers may therefore play a major role in determining patterns of gene flow across Australia.

Interestingly, while starlings in the native Palearctic and invasive North American ranges display migratory behaviour, this has been lost in Australian starlings (Cabe 1999, Rollins, *et al*. 2009). While starlings may move in excess of 100’s of kms from their natal site (Feare 1984), a lack of largescale Australia-wide interbreeding is evident in their underlying genetic population structure. Previous genetic surveys based on microsatellite markers showed that starlings in Australia form four distinct genetic clusters (Rollins, et al. 2009). The genetic cluster representing the range-edge at the western most extent of the starlings’ range in Western Australia (WA) contained novel genetic variants relative to range-core and native individuals, and evidence of genetic drift and selection was identified in this cluster (Rollins, et al. 2016). This stands in contrast to starling populations of North America, also introduced in the late 19^th^ Century, for which high dispersal and continental wide interbreeding appears to have led to genetic homogenisation, minimal population substructure, and lower F_ST_ values between sites, when compared to the Australian starlings (Cabe 1998, 1999). The Australian invasion is characterised by lower genetic diversity than that of the native range (Rollins, et al. 2011). Nevertheless, there is clear evidence of morphological variation in response to temperature and rainfall gradients across the invasive range in Australia (Cardilini, et al. 2016).

Given the wide range of environments that Australian starlings inhabit and the approximately 60 generations since introduction, the invasive starling provides an excellent model for investigating the population dynamics and adaptive variation of a highly dispersive invasive species. Further, the multiple Australian introductions enables us to examine whether introduction regime may leave genetic footprints in modern day populations. Here, we use a reduced representation sequencing approach to generate a SNP data set to assess the roles the Australian landscape and starlings’ invasion history have played in shaping the genetic structure and patterns of adaptive genetic variation found across the Australian range. We aim to test two non-mutually exclusive hypothesis: that genetic differentiation will be related to geographic distance, and that genetic differentiation will be related to environment gradients. To do this, we compare population structure to underlying environmental differences and geographic distances separating sites. Finally, we then use three different analyses to test for adaptive genetic variation across the starlings geographic range in Australia.

## Methods

### Sample collection and sexing

We collected a total of 568 starling samples from 24 localities across the invasive range in Australia (Fig. 1b, Table 1). Sample collection occurred in two separate phases. First, we used a subset of pre-extracted DNA samples (n = 150) collected by Rollins *et al*. (2009), from localities across South Australia and Western Australia from 2003 – 2007. Second, we collected 418 samples between May 2011 and October 2012 from 18 locations across the south-eastern portion of the Australian range (Fig. 1b). These birds were collected in three ways: *i)* by trapping, *ii)* as carcasses collected from 3^rd^ party hunters or landowners that humanely killed birds on their property, or *iii)* chicks collected directly from nests. In the latter case, only one individual per family (adult or chick) was included in the dataset to avoid the inclusion of closely related individuals. We recorded Global Position System (GPS) coordinates upon collection. When 3^rd^ party hunters did not record GPS coordinates, the closest corresponding address was recorded and GPS coordinates were identified using Google maps. We took a muscle sample from the thigh or breast and stored it in 70% ethanol prior to DNA extraction. We extracted DNA with a Gentra PureGene Tissue Kit, following the manufacturer’s protocol. All samples were sexed using molecular methods (Griffiths, et al. 1998).

**Table 1.** Summary statistics for each starling collection locality. N = number of samples collected from each location, GC = the genetic cluster from whole-genome dataset (1 = Munglinup cluster, 2 = southern Australia cluster, 3 = southwest New South Wales cluster, 4 = northern Australia cluster), OGC = the genetic cluster from outlier dataset (1 = Western and Southern Australia, 2 = Eastern and Northern Australia), H_O_ = Observed Heterozygosity, H_S_ = the within population gene diversity, Locality F_ST_ = fixation index for each population against all other individuals, % Poly SNPs = the percentage of polymorphic SNPs. We do not report a Tajima’s D value for Streaky Bay due to low sample size.

| Collection Locality | Sample site number | State | N | GC | OGC | Env. Cluster | $H_O$ | $H_S$ | Locality $F_{ST}$ | % Poly. SNPs | Tajima's D |
| --- | --- | --- | --- | --- | --- | --- | --- | --- | --- | --- | --- |
| Munglinup | 1 | WA | 18 | 1 | 1 | 1 | 0.092 | 0.225 | 0.094 | 55.2 | 0.137 |
| Condingup | 2 | WA | 16 | 2 | 1 | 1 | 0.092 | 0.257 | 0.051 | 66.1 | -0.129 |
| Coorabie | 3 | SA | 19 | 2 | 1 | 1 | 0.137 | 0.265 | 0.028 | 80.9 | -0.395 |
| Streaky Bay | 4 | SA | 9 | 2 | 1 | 1 | 0.069 | 0.267 | 0.137 | 35.9 | NA |
| Tumby Bay | 5 | SA | 22 | 2 | 1 | 1 | 0.114 | 0.269 | 0.030 | 83.3 | -0.134 |
| McLarenavale | 6 | SA | 20 | 2 | 1 | 1 | 0.141 | 0.266 | 0.032 | 86.0 | -0.423 |
| Warrnambool | 7 | VIC | 24 | 2 | 1 | 3 | 0.117 | 0.271 | 0.026 | 88.6 | -0.286 |
| Bendigo | 8 | VIC | 25 | 2 | 1 | 2 | 0.159 | 0.273 | 0.027 | 93.4 | -0.449 |
| Wonthaggi | 9 | VIC | 22 | 2 | 1 | 3 | 0.119 | 0.272 | 0.030 | 81.6 | -0.292 |
| Orbost | 10 | VIC | 22 | 2 | 1 | 3 | 0.148 | 0.270 | 0.028 | 89.0 | -0.468 |
| King Island | 11 | TAS | 14 | 2 | 1 | 3 | 0.135 | 0.272 | 0.027 | 80.9 | -0.395 |
| Sheffield | 12 | TAS | 21 | 2 | 1 | 3 | 0.098 | 0.265 | 0.037 | 79.2 | -0.106 |
| Cygnets | 13 | TAS | 17 | 2 | 1 | 3 | 0.095 | 0.262 | 0.037 | 72.5 | -0.190 |
| Hay | 14 | NSW | 26 | 3 | 1 | 2 | 0.158 | 0.259 | 0.025 | 86.2 | -0.581 |
| Albury | 15 | NSW | 24 | 3 | 1 | 2 | 0.200 | 0.259 | 0.038 | 89.5 | -0.899 |
| Nowra | 16 | NSW | 25 | 4 | 2 | 3 | 0.133 | 0.268 | 0.030 | 89.2 | -0.278 |
| Maitland | 17 | NSW | 25 | 4 | 2 | 3 | 0.149 | 0.266 | 0.034 | 91.7 | -0.391 |
| Dubbo | 18 | NSW | 20 | 3 | 2 | 2 | 0.165 | 0.257 | 0.030 | 82.9 | -0.671 |
| Nyngan | 19 | NSW | 25 | 4 | 2 | 2 | 0.126 | 0.273 | 0.023 | 84.7 | -0.391 |
| Austral Eden | 20 | NSW | 21 | 4 | 2 | 3 | 0.142 | 0.273 | 0.024 | 87.0 | -0.349 |
| Tamworth | 21 | NSW | 11 | 4 | 2 | 2 | 0.129 | 0.271 | 0.029 | 64.6 | -1.409 |
| Moree | 22 | NSW | 24 | 4 | 2 | 2 | 0.166 | 0.271 | 0.028 | 91.6 | -0.546 |
| Lismore | 23 | NSW | 22 | 4 | 2 | 3 | 0.123 | 0.270 | 0.024 | 86.9 | -0.393 |
| Lemon Tree | 24 | QLD | 24 | 4 | 2 | 2 | 0.141 | 0.272 | 0.024 | 89.0 | -0.418 |

### Environmental data collection

For each collection locality, we extracted climatic variables from Bioclim data sets using the raster package in R (Hijmans 2016). Climatic variables included: mean diurnal range (Bio02), isothermality (seasonal temperature variation) (Bio03), temperature seasonality (Bio04), maximum temperature of the warmest month (Bio05), minimum temperature of the coldest month (Bio06), temperature annual range (Bio07), precipitation of the wettest month (Bio13), precipitation of the driest month (Bio14) and precipitation seasonality (Bio15). We extracted the aridity index (AI) data from the CGIAR-CSI Global-Aridity and Global-PET Geospatial Database (Zorner, et al. 2008). We downloaded monthly average normalised difference vegetation index (NDVI) values from the Australian Bureau of Meteorology for the period 1997-2011. We averaged monthly datasets across years (e.g. mean NDVI January 1997-2011) and then across all months to create a mean NDVI raster file from which we extracted mean NDVI (mnNDVI) for each sample. We calculated variability in day length (varDL) per sample using the ‘daylength()’ function in the R package geosphere (Hijmans, et al. 2015). We defined the position of each collection locality as the central longitude and latitude of all samples collected from the same collection locality. Using an R function that we wrote, we extracted elevation data (Elev) for each sample collection locality from Google maps (see supplementary file for function). We calculated the average of each environmental variable at each collection locality and used these for further analysis. Using the R package sp (Bivand, et al. 2013), we calculated geographic distance variables as Euclidean distance (km) of each locality to the nearest coastline and Euclidean distance of each locality to nearest known site of introduction. To identify a reduced set of variables that explained environmental variation across the introduced range, we conducted a principal components analysis (PCA) including all environmental variables and run in R using the prcomp function (RCoreTeam 2015). We calculated the pairwise distance between points on a plot of PC1 and PC2 to calculate pairwise environmental distances between collection localities, which were subsequently used in the analysis of isolation by environment (IBE matrix) (Wang 2013; Wang, et al. 2013; Leydet, et al. 2018). Finally, using the R package sp (Bivand, et al. 2013) we calculated pairwise distances as the greater circle distances between collection localities to test for isolation by distance (IBD matrix).

These environmental data were used to examine IBE, IBD, and the relationship between geographic and environmental distances across sampling sites (see “Genetic, environmental and geographic relationships”), and in the identification of candidate loci under selection (see “Detecting putative loci under selection”).

### Library construction, sequencing and SNP calling

The Cornell University Institute of Biotechnology Genomics Facility conducted library construction and Genotyping-by-Sequencing (GBS) (Elshire, et al. 2011), using the restriction enzyme *PstI*. Each individual received a unique barcode before multiplexing 96 individuals per lane (Illumina HiSeq 2000; 6 lanes, 100 bp single-end reads).

We processed raw sequence data using TASSEL 3.0 and called SNPs for all individuals using the UNEAK pipeline (Bradbury, et al. 2007; Lu, et al. 2013; Glaubitz, et al. 2014). We used default values for parameters and sequence read (tags) that were detected at least five times across all samples were retained. We compared tags (aligned 64 bp reads) using network filtering (identifies tag pairs by filtering out repeats, sequencing errors, and paralogs) and considered those with a 1 bp mismatch to be candidate SNPs, using an error tolerance rate of 0.03 (Lu, et al. 2013). After these initial filtering steps, 6,102,279 reads remained and we combined and converted into Tags on Physical Map format (creating a pseudo chromosome), which acted as a reference. SNPs were called for each individual and we identified a total of 291,189 SNP loci. When calling SNPs against the reference pseudochromosome, we only considered tags recorded in at least three individuals. We used a maximum genotype depth of 60 (~10X the average genotype depth of the dataset) for all data sets to filter out genotypes that were possibly overrepresented (due to being in repetitive genomic regions) (Almeida, et al. 2014). We removed any locus that was missing in more than 90% of samples and, subsequently, removed all samples that had less than 10% of the total number of loci. We tested our data for conformity to Hardy Weinberg Equilibrium (HWE) (VCFTools, v0.1.12b) (Danecek, et al. 2011) and, after correcting *p*-values for multiple comparisons using FDR 0.05 (Benjamini and Hochberg 1995), only loci identified out of HWE in five or more collection localities were to be removed; no loci were found to be out of HWE in five or more collection localities.

We examined a range of filtering metrics (detailed methods in Supplementary Material, Fig. S1) and chose a final dataset with a minor allele frequency (MAF) of 0.05, a minimum genotype depth of three and maximum level of missingness of 50%, resulting in the inclusion of 16,177 SNP loci across 499 individuals (data accessible at https://github.com/katarinastuart/Sv1_StarlingGBS/tree/master/Data). We chose this dataset because including tags with large proportions of missing data can obscure signal, while stringent missingness filtration by definition reduces diversity, leading to an underestimation of population structure (Huang and Knowles 2016). Regarding MAF, higher thresholds are proven to inhibit population substructure detection, a problem that also persists when singletons are retained (i.e. low MAF threshold) (Linck and Battey 2019). Hereafter this SNP dataset will be called the genome-wide dataset.

### Population structure analysis

#### Population structure based on genome-wide SNP loci

Global F_ST_ and pairwise F_ST_ values were estimated in the R package hierfstat (Goudet 2005), as were observed heterozygosity [H_O_] and within population gene diversity [H_S_]. Upper and lower confidence intervals (95%) were obtained for F_ST_ from 10,000 bootstraps, *p*-values were corrected for multiple comparisons using FDR 0.05 (Benjamini and Hochberg 1995). Collection locality F_ST_ (F_ST_ of one population against all other populations) and Tajima’s D (Tajima 1989) values within all collection localities were calculated in the R package popgenome (Pfeifer, et al. 2014).

We determined the number of distinct genetic groups in Australia using a Bayesian assignment test in the program fastSTRUCTURE (v1.0) (Raj, et al. 2014). Additionally, we analysed population structure of a neutral data set, to ensure it was reasonable to use the genome-wide dataset without removing SNPs contributing to non-neutral variation (detailed methods in Supplementary Material).

#### Population structure based on outlier SNP loci

We identified F_ST_ outlier loci in Bayescan (v2.1) (Fischer, et al. 2011), which uses a Bayesian approach to compare the F_ST_ values of a locus across subpopulations (Foll and Gaggiotti 2008). Bayescan was run for two different groupings of the samples. on the first run, we allocated individuals to one of 24 subpopulations that related to their collection localities. Bayescan is prone to high levels of false positives; however, Lotterhos and Whitlock (2014) showed that this effect can be reduced by increasing the prior odds value. Bayescan was run with default parameters except that the prior odds value was set to 100. Only loci with a Bayes factor greater than three were kept as outliers because values above this indicate substantial evidence of selection (Jeffreys 1961). We determined the number of distinct genetic groups determined by the outlier loci dataset identified with BayeScan using fastSTRUCTURE (detailed methods in Supplementary Material). Additionally, to validate the outlier loci data set, we generated a random subset of an equivalent number of SNPs from the genome-wide dataset and analysed these as above (detailed methods in Supplementary Material).

### Genetic, environmental and geographic relationships

We used Multiple Matrix Regression with Randomization analysis (MMRR) (Dryad doi:10.5061/dryad.kt71r) to determine if geographic distance between localities predicts environmental distance. Then, we used MMRR analysis to determine if geographic distance between localities (IBD matrix) predicts genetic distance (isolation by distance, IBD), or if environmental distance (IBE matrix) predicts genetic distance (isolation by environment; IBE). MMRR was implemented by comparing a pairwise genetic distance matrix with IBD and IBE distance matrices following Wang (2013). Briefly, random permutations are conducted between the rows and columns of the dependant (genetic distance) matrix to produce the resulting matrix and its coefficients (while other matrices remain fixed) - a null distribution is formed from this permutation, and observed values may be compared to it to determine if there is any relation between matrix distances (Legendre, et al. 1994).

### Detecting putative loci under selection

We investigated adaptive variation across the starling’s range using candidate loci identified by Bayscan (F_ST_ outlier loci) as outlined above, and through two different methods of environmental association analyses (Bayenv2 and RDA). Bayescan was run with as above.

While measures of population differentiation such as F_ST_ allow for SNP’s putatively under selection to be identified, measured differentiation may simply be a result of neutral variation (demographic effects, drift etc.), and such an approach does not directly allow possible selective drivers to be identified (e.g. environment) (Feng, et al. 2015). Hence it is common for scans for selection to also include environmental association analysis (Pluess, et al. 2016), such as logistic regression. The nature of adaptive drivers though means a diverse array of factors may be subtly influencing genomic sites, so many regressions may need to be performed if there are a number of environmental drivers of interest. To address this, constrained ordination methods are fast becoming a popular method when conducting genotype-environment associations with multivariate and multilocus datasets, boasting detection power coupled with low false-positive and high true-positive rates (Forester, et al. 2018), and therefore are invaluable for investigating adaptation (Capblancq, et al. 2018). Under the conditions of genotype-environment associations, constrained ordination techniques such as RDA represent parsimonious models that attempt to capture total population variation specifically explained by constraining predictor (e.g. environmental) variables. Thus, RDA environmental association analysis was conducted alongside the traditional allele frequency association tests. However, RDA cannot be conducted on matrices with missing data, so imputation on missing values is necessary – a particular problem for dataset like GBS that are characterised by high levels of missingness. It is possible that imputation might lead to type I and type II errors, hence we used this method alongside the two more traditional analyses (Bayscan and Bayenv2) to test for signals of selection.

We used Bayenv2 (v2.0; Gunther and Coop 2013) to test for association of minor allele frequencies with environmental variables. Bayenv2 calculates the correlation between allele frequencies and environmental variables and was used to tests for associations between all SNPs and three environmental variables (Coop, et al. 2010; Gunther and Coop 2013). The three environmental variables that were tested included Aridity, Bio05 (maximum temperature of the warmest month), and Bio 15 (precipitation seasonality). These variables were chosen because temperature and precipitation have previously been shown to influence starling phenotype (Cardilini, et al. 2016). We estimated the covariate matrix using a random subset of 2,000 loci. Runs were set at 100,000 iterations and we kept any locus with a Bayes factor greater than three as being potentially under selection (Gunther and Coop 2018). From the candidate SNPs identified by BayEnv2, we extracted the SNP that most highly correlated with each of Aridity, Bio05, and Bio 15, and plotted allele frequency against the environmental variables score.

We used the R package vegan (v3) (Oksanen, et al. 2018) to conduct redundancy analysis (RDA) to determine if any particular SNP locus is heavily loaded onto predictor axes, from which we may infer the occurrence of selection. The genome-wide SNP dataset was loaded against the climatic variables associated with our sampling sites. The SNP genotype dataset was processed into 012 format (count reflects the number of the minor allele) using VCFtools, and missing SNP genotypes (average missingness across all sites: 44.3%) were imputed based on probabilities for each given genotype at that genome locality using a custom script (detailed methods in Supplementary Material). To reduce multicollinearity, from the complete list of environmental variables, we retained seven predictors with relatively low variance inflation factors (range: 1.9—6.3) to reduce multicollinearity: (Elev, Bio03 (isothermality); Bio06 (minimum temperature of the coldest month); Bio13 (precipitation of the wettest month); Bio15 (precipitation seasonality); mnNDVI; and varDL. We used the function anova.cca in vegan to confirm the model’s significance. Once the SNP data were loaded against the RDA axes, candidates for selection were determined to be those that lay more than three standard deviations away from the mean (Forester, et al. 2018).

We extracted the associated 64 bp tags for loci identified by at least one of the methods of Bayescan, Bayenv2, and RDA. These tags were searched against the annotated starling genome available on NCBI (GCF_001447265.1) and functional descriptions extracted, following the protocol described in Richardson and Sherman (Richardson and Sherman 2015). Briefly, 64 bp tags of outlier loci were searched against the starling predicted RNAs using Blastn with a e-value cutoff of 1.0×10^-3^ (Camacho, et al. 2009). Gene names were extracted from the blast output for the best hit matches per outlier loci (smallest e-value, highest % identity and length match) and these gene names were matched to Uniprot IDs and associated functional descriptions extracted (Bairoch, et al. 2008).

## Results

### Population environmental distribution

A PCA analysis of environmental parameters grouped collection localities into three distinct clusters (Fig. 1b; Supplementary Material, Fig. S2, Table S1). These clusters defined three distinct environmental regions: (i) Western Australian and South Australian collection localities (arid, blue); (ii) eastern Australia collection localities inland of The Great Dividing Range (semi-arid, red); and (iii) collection localities on the coastal side of The Great Dividing Range (non-arid, green) (Supplementary Material, Table. S1).

### Population structure analysis

#### Population structure based on genome-wide SNP loci

Analysis of the patterns of genetic diversity revealed that populations at the expansion front in Western Australia (Munglinup and Condingup) displayed relatively lower levels of genetic diversity compared to populations in the east of the introduced range (Table 1). Further, Munglinup was the only population to have a positive Tajima’s D (low frequency of rare alleles). The global level of genetic differentiation across collection localities was low, but significant with an average F_ST_ of 0.027 (95% CI 0.025 - 0.030) (Supplementary Material, Table S2). Pairwise comparisons of F_ST_ between collection locations revealed significant genetic structuring between 263 out of 276 pairwise comparisons (Supplementary Material, Table S2). Collection locality (location vs all others) pairwise F_ST_ values revealed that those populations with the lowest levels of genetic diversity also showed the greatest level of genetic differentiation to all other populations (Table 1).

The Bayesian structure analysis based on the full dataset of 16,177 SNPs showed that K=2, and K=4 had similar levels of support (Supplementary Material, Fig. S3a), while K=4 had the most support when considering only neutral genome-wide loci (Supplementary Material, Fig. S3a). For the former, the four genetic clusters were Munglinup, southern Australia, southwest New South Wales, and northern Australia (Fig. 1b, Fig. 2). Within the fastSTRUCTURE plot, the Munglinup and southwest New South Wales clusters were relatively distinct though admixture is still present, while the two larger genetic clusters were less well defined suggesting more admixture (Fig 3). Subsequent fastSTRUCTURE analysis found no substructure within any of the four genetic clusters, and analysis on the neutral dataset produced a very similar likelihood plot (Supplementary Material, Fig. S3b).

**Fig. 2.**
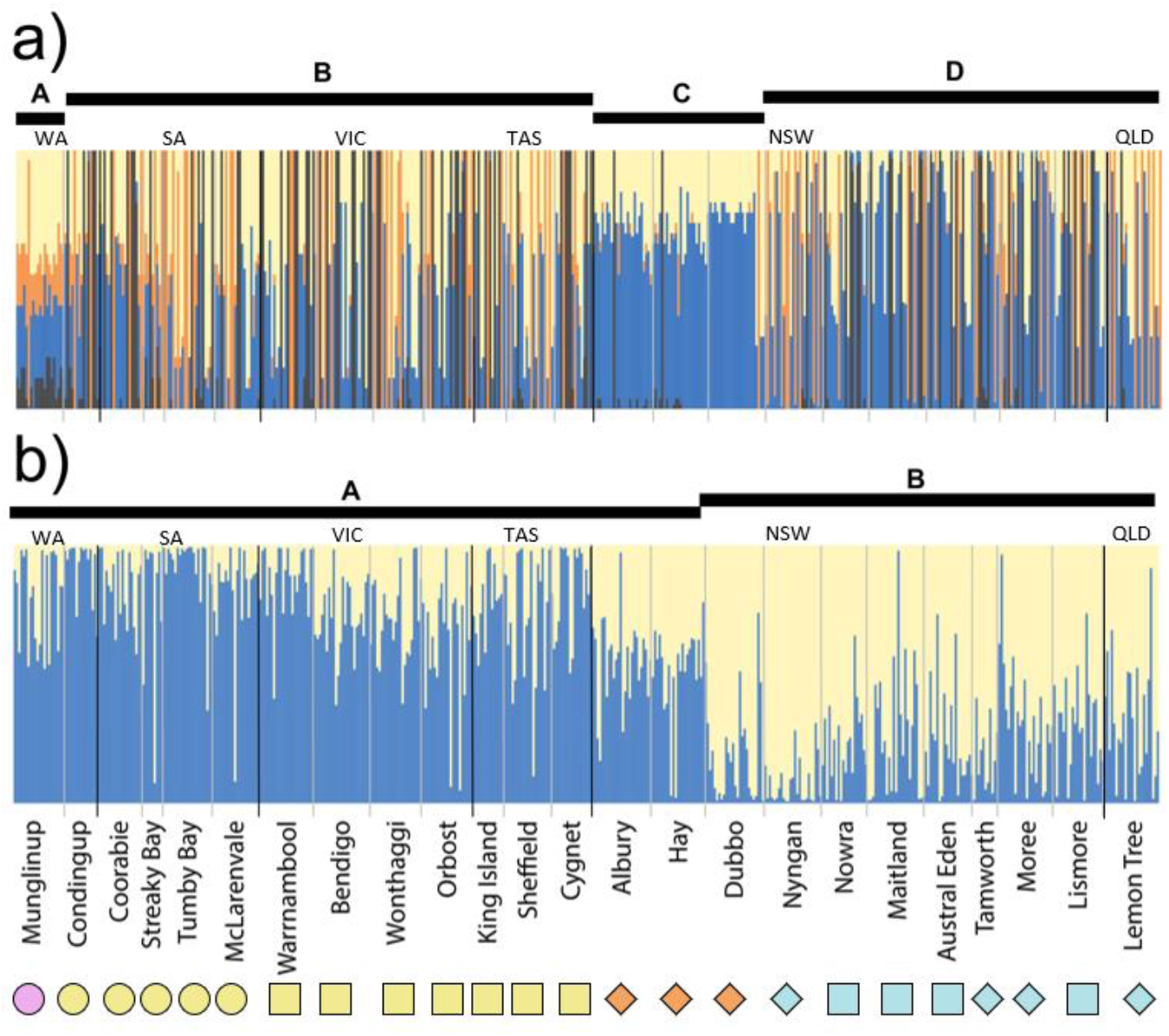
fastSTRUCTURE plots indicating the genetic structure of Australian starling populations. Each column represents an individual sample, with the colours indicating the estimated likelihood that the individual belongs to a particular genetic group. Collection localities are arranged from west to northeast following geographic distribution. The vertical black lines demarcate state political boundaries indicated in the abbreviations above each plot, and the vertical grey lines delimit collection localities. The black bars at the top of the graph and their corresponding names identify genetic clusters. Each panel is derived from a different dataset with a) indicating genetic structure of the genome-wide dataset when K = 4 and b) indicating the genetic structure of the outlier dataset when K = 2. Labels indicate the distinct genetic clusters reported by the fastSTRUCTURE analysis for each panel; panel a) Munglinup (A), southern Australia (B), southwest New South Wales (C), northern Australia (D); panel b) southern Australia (A), northern Australia (B). Symbols and colour used at the bottom of the figure are based on Fig. 1b.

#### Population structure based on outlier SNP loci

Out of the 16,177 SNPs, Bayescan analyses identified 89 outlier loci (Table 2) as candidates for selection (log10(BF) > 3), while all other loci were considered neutral. These loci made up the outlier data set. As expected, pairwise F_ST_ values calculated for outlier loci were considerably higher than those calculated for the genome-wide dataset, with an average F_ST_ of 0.122 (95% CI 0.101 – 0.130, non-overlapping with the CI for genome-wide SNP loci calculated F_ST_) and most locality F_ST_ comparisons for the outlier dataset were significant (256/276; Supplementary Material, Table S2). fastSTRUCTURE analyses of the outlier dataset identified two distinct genetic clusters which covered the larger clusters from the genome wide SNP dataset, with the range edge cluster integrated into the eastern cluster, and the inner NSW cluster split amongst the two new large outlier clusters (Fig. 2; Supplementary Material, Fig. S3c), whereas we found no genetic structure in the random subset of loci (Supplementary Material, Fig. S3d). Subsequent fastSTRUCTURE analysis found no substructure in the two main genetic clusters (data not presented).

**Table 2.**
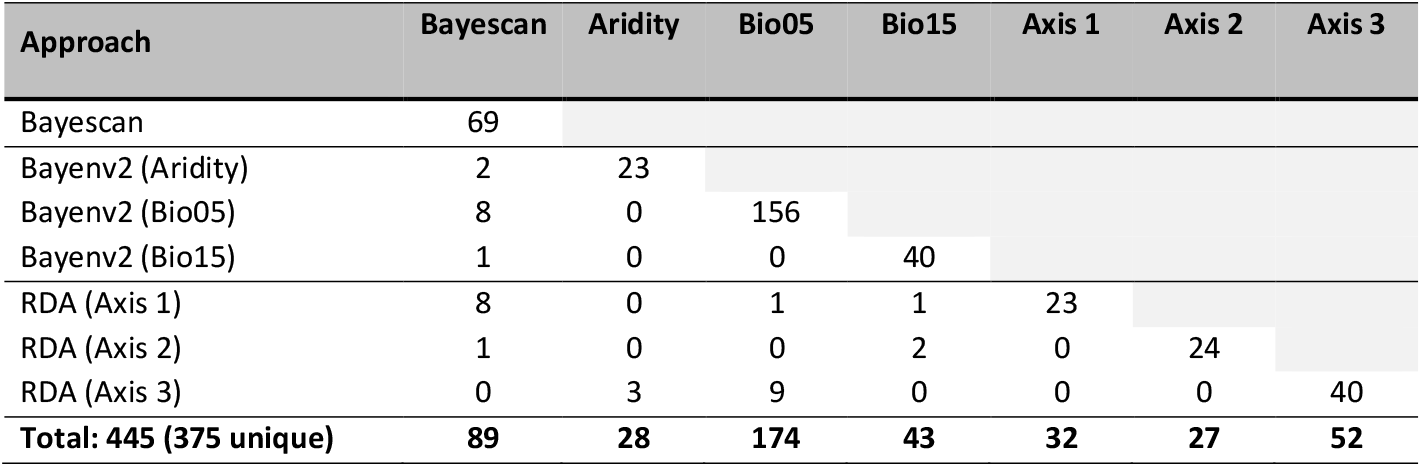
Summary of candidate SNPs under putative selection, as detected by Fst outlier (Bayescan) and environmental association (Bayenv2 and RDA) approaches. Bayenv2 analysis is broken down into the three tested environmental variables Aridity, Bio05 (maximum temperature of the warmest month), and Bio 15 (precipitation seasonality). RDA is broken down into three key RDA axes. Displayed are number of unique SNPs identified for each method (diagonal), total number of SNPs identified for each method (bottom line) and SNPs common to each pairwise comparison of approaches (bottom matrix).

### Genetic, environmental and geographic relationships

Geographic distance did significantly correlate with environmental distance across the starling’s Australian range (Fig. 3), so IBE and IBD were calculated in the same MMRR model to account for confounding measures of geographic distance and environmental distance. When genetic distance was tested against geographic distance and environmental distance in an MMRR, only IBD was found to explain any variation in genetic differentiation observed between collection localities (Fig. 4a). IBE was not found to be a significant predictor of genetic distance (Fig. 3). When the genetic distance of the outlier dataset was tested against geographic distance and environmental distance in an MMRR, only IBD was found to explain any variation between collection localities (Fig. 4b). IBE was not found to be a significant predictor of genetic distance (Fig. 3).

**Fig. 3.**
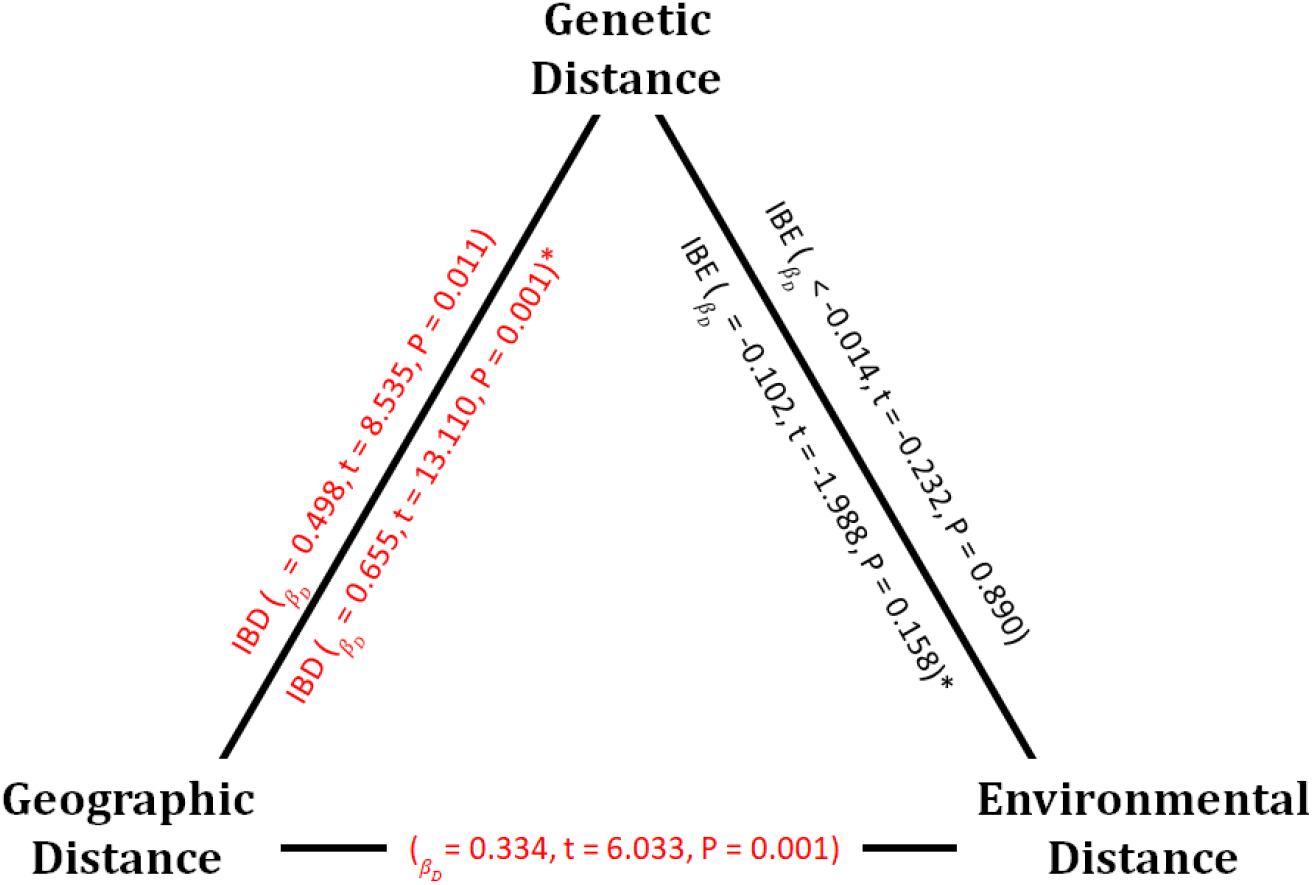
Results of Multiple Matrix Regression with Randomization analysis (MMRR) capturing genetic, environmental and geographic relationships in Australian starlings. Analysis results on the outside of the triangle were performed on the genome-wide dataset (r^2^ = 0.229), while analysis results listed inside the triangle and with an asterix (*) indicates analysis that were performed on outlier loci data set (r^2^ = 0.396). Analysis results of geographic distance and environmental distance is not specific to a genetic data set (r^2^ = 0.117). Results coloured red indicate significant results.

**Fig. 4.**
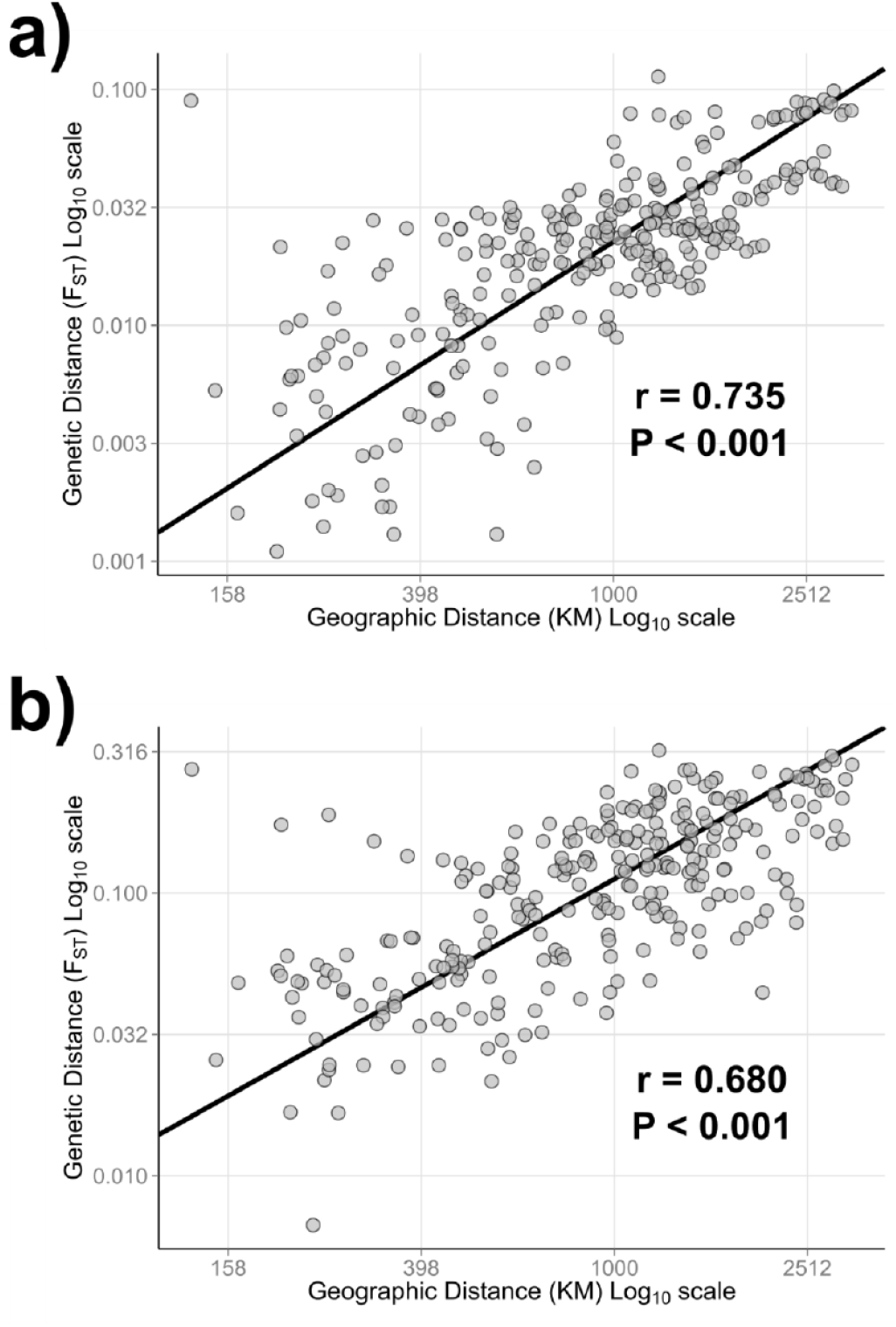
Plots of isolation by distance (IBD) in Australian starlings with Mantel correlation and p-value indicated. Each point represents a pairwise relationship between collection localities and the line represents the linear relationship between genetic distance and geographic distance. Plot a) represents IBD of the full dataset, while plot b) represents IBD of the outlier dataset. Both genetic distance and geographic distance were log_10_ transformed.

### Detecting putative loci under selection

Using three different methods to identify SNPs putatively under selection, we found a total of 375 different SNPs. Bayescan analyses of F_ST_ identified 89 outlier loci (Table 2). BayEnv2 identified positive associations between 245 SNPs and three environmental variables including Aridity (N = 28 SNPs), Bio05 (temperature) (N = 174 SNPs), and Bio15 (precipitation) (N = 43 SNPs) (Table 2). The minor allele frequency of the locus most strongly associated with Aridity (ID = 68063014) showed a negative relationship (C = −0.272, SE = 0.070, t = −3.883, P < 0.001, r^2^ = 0.380; Fig. 5a). The minor allele frequencies of the SNP (ID = 233190026) most strongly associated with Bio05 showed a strong positive relationship (C = 0.018, SE = 0.004, t = 4.116, P < 0.001, r^2^ = 0.409; Fig. 5b). The relationship between Bio05 and the minor allele frequency for this SNP is closely aligned with geography, with the Munglinup cluster and almost all collection localities from the southern Australia cluster being fixed at this locus. Loci associated with Bio15 (ID = 196163030) showed a positive relationship (C = 0.008, SE = 0.002, t = 3.607, P = 0.002, r^2^ = 0.343; Fig. 5c), with most individuals from the northern Australia cluster being fixed at this locus.

**Fig. 5.**
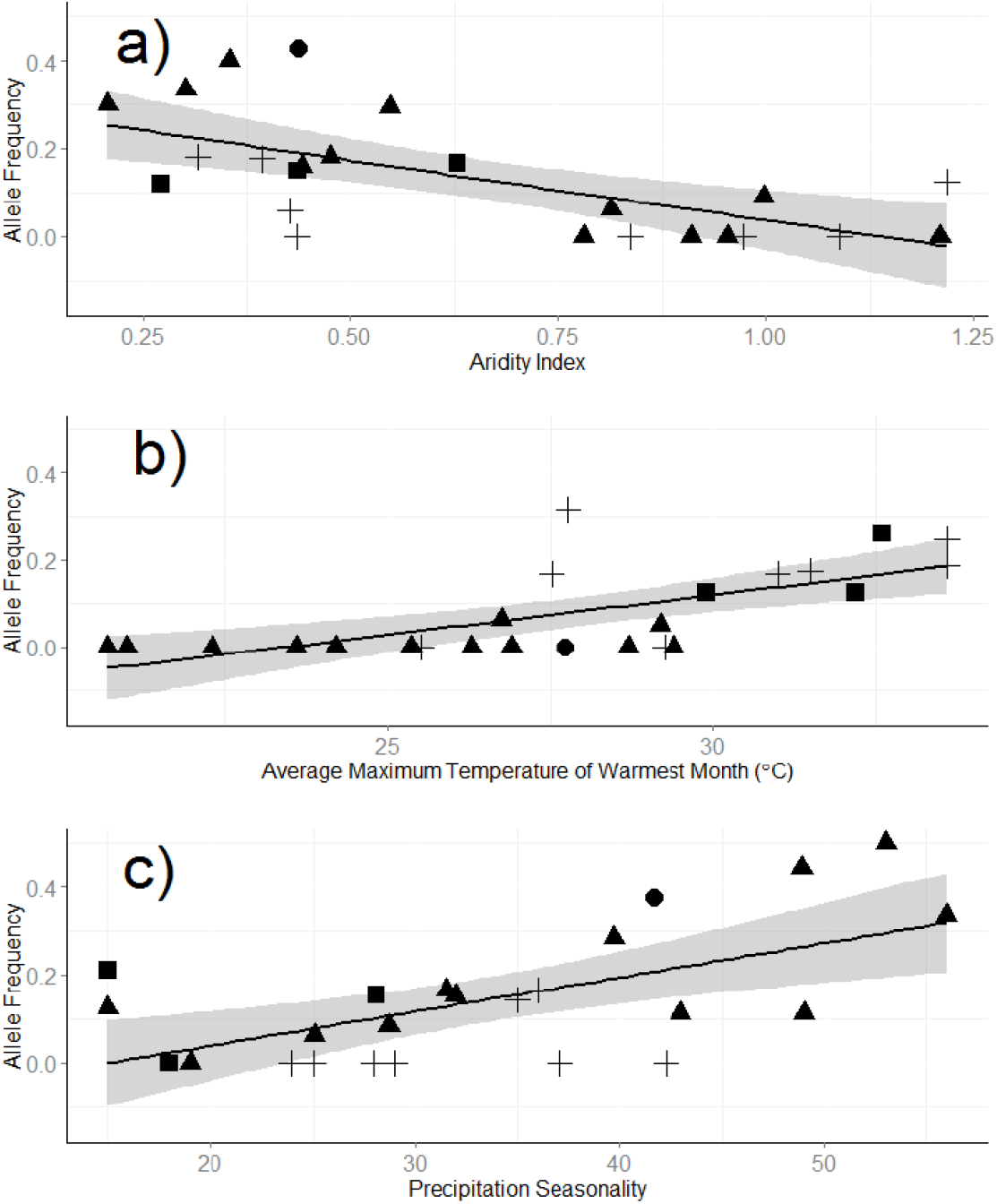
The relationship between starling collection locality allele frequency and a locus strongly associated with a) Aridity (SNP ID 68063014), b) Bio05 (SNP ID 233190026), and c) Bio15 (SNP ID 196163030). The black line in each panel shows the linear relationship between the environmental variable and allele frequency, with the grey ribbon indicating the 95% confidence interval. Each point represents a collection locality with its shape indicating the genetic cluster that it belonged to; circle = Munglinup cluster, triangle = southern Australia cluster, square = southwest NSW, and cross = northern Australia cluster.

The RDA model was significant (F_7,491_ = 1.2228, P < 0.001; Fig. 6), and reported population clustering largely concurring with the results of the PCA environmental parameters (Supplementary Material, Fig. S2). RDA identified 111 candidate SNPs that showed strong association with the local environment (first 3 axes reported significance and were used for SNP identification; F_1,491_ = 2.0516, P < 0.001; F_1,491_ = 1.2855, P < 0.001; F_1,491_ = 1.1708, P < 0.008). The genetic variation between the westernmost starlings (WA and SA) and NSW starlings correlated with a number of variables: Bio03 (Isothermality); Bio06 (Min Temperature of Coldest Month); Bio15 (Precipitation Seasonality); Elev (Elevation); mnNDVI (mean vegetation cover) (Fig. 6). Meanwhile, genotype differences between southern starlings (VIC and TAS), and northern starlings (NSW and QLD) correlated with varDL (Variability in Day Length) and Bio13 (Precipitation of Wettest Month) (Fig. 6).

**Fig. 6:**
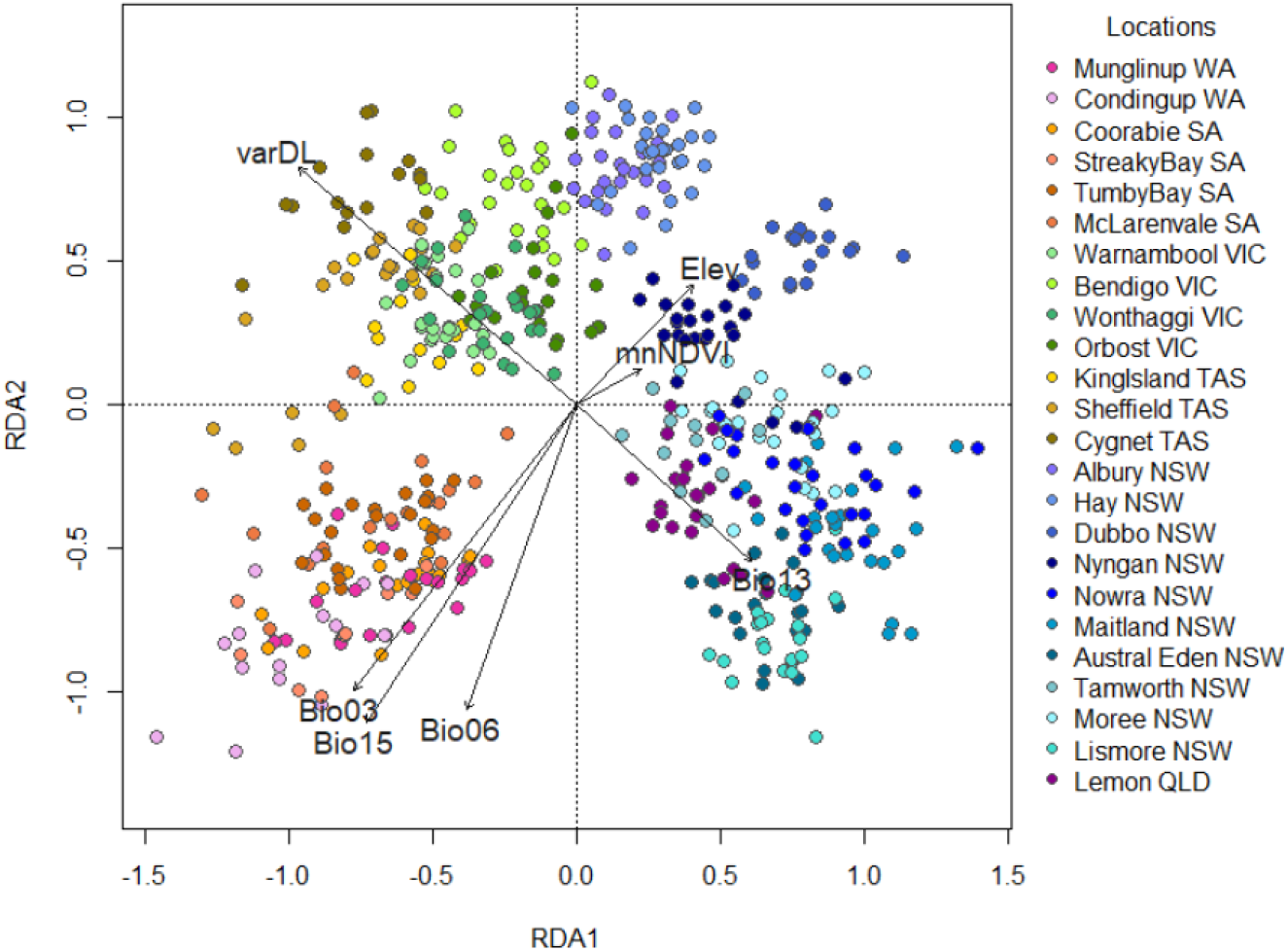
The ordination plot of redundancy analysis (RDA) of 16177 SNP data for 499 starlings, collected from 24 sample sites across 5 Australian states. Individuals are coloured by location, black arrows represent selected environmental variables. The relative placement of environmental predictor arrows to individual and location sample clustering represents the correlation between that environmental variable and the genetic differences seen in that cluster.

From the 375 unique SNPs identified across all three methods (Table 2), a total of 25 proteins were identified when mapped to the starling reference genome (Supplementary Material, Table S3). Of these 25 mapped proteins, 7 loci were associated with Bayescan F_ST_ outliers, 11 were associated with Bayenv2 environmental variables, and 8 were associated with candidate loci identified through the RDA environmental association (Supplementary Material, Table S3). Among the annotated SNPs, we see a range of biological functions traditionally associated with adaptation in invasive species, including processes such as immune system responses (*C1QC* and *STAB1*) (Colautti, et al. 2004), temperature tolerance (*HSPA9*), stress (*TRAF1*) (Zerebecki and Sorte 2011), and dietary alterations (*CTRL*) (Spit, et al. 2014).

## Discussion

This study found that since their introduction into Australia, starling populations have become genetically differentiated despite the potential for high levels of dispersal, and that selection has facilitated their adaptation to the wide range of environmental conditions across their geographic range. Isolation by distance appears to have played a strong role in determining genetic substructure across the starling’s Australian range, but neutral population structure does not seem to be affected by environmental factors (IBE). Analyses of candidate SNPs that are putatively under selection indicate that aridity, precipitation, and temperature may be important factors driving selective change across the starling’s invasive range in Australia, indicating genetic differentiation in response to environment may be occurring at a limited number of loci. We discuss some of our findings alongside a similar study on the invasive North American starlings and explore population similarities and differences across these replicated invasions. We do note that some loci that are candidates for selection suggest that the faint genetic footprint left by the historic regime may hide, or be mistaken as, putative adaptation.

### Population structure

#### Population structure of Australian starlings

Our genome-wide SNP data set identified between two and four distinct genetic clusters. When the genome wide data set was analysed at K=4, two of the genetic clusters covered large geographic regions; the first encompassing southern Australia (WA to TAS), and the other encompassing northern Australia (NSW and southern QLD). Munglinup and arid inland NSW represented two small distinct clusters, indicating higher population isolation in these areas. The population genetic structure of Australian starlings identified with the genome-wide dataset of SNPs was similar to that identified in previous work, but contained some key differences (Rollins, et al. 2009; Rollins, et al. 2011). Microsatellite data support the presence of four populations: two small, distinct incursions into WA and two large populations, one in SA, and another including VIC, TAS, and NSW (Rollins et al. 2009). It is likely that microsatellite variation is less sensitive given the reduced power of the data set to characterise differences, when compared to the SNP data set.

Interestingly, Munglinup represented the only population that reported a positive Tajima’s D (Table 1), which suggests demographic (e.g. a sudden population contraction) or selection effects. Munglinup represents the western-most range edge, and it was likely established from a small founding number of individuals with limited on-going gene flow and potential selection, thereby providing conditions for more rapid divergence. Despite the fact that Munglinup is only approximately 150km away from the next sampling site (Condingup), we see distinct genetic differences at the range edge. Similar results were found in previous studies (Rollins et al. 2011), which support the idea that the Munglinup subpopulation is a result of a later, separate incursion into WA, and not a result of dispersion of Condingup individuals, demonstrating that expanding ranges edges may be the site of complex demographic processes.

Population structure analysis using the outlier loci dataset revealed only two population clusters. A restricted outlier dataset (though prone to false positives) may allow for the detection of adaptive variation, and has been used in many prior studies to further resolve population structure than just using neutral loci (Hess, et al. 2013; Milano, et al. 2014; Benestan, et al. 2015). In this study, the genome-wide dataset identifying four genetic clusters while the outlier loci identified two genetic clusters (Fig. 2). What the results may indicate is two main genetic clusters may be differentiated by adaptive differences, whereas demographic effects may be responsible for the structure seen using neutral loci.

The two large genetic clusters identified in the genome-wide and outlier loci dataset largely overlap, and may be explained by two competing phenomena: 1) a combination of geographic isolation and/or historic genetic diversity from different source populations, or 2) adaptive selection driving genetic differentiation. Strong IBD, and the fact that the higher subpopulation structure resolution of the genome-wide dataset determined that subpopulations were not centred around unique introduction sites, points to the latter explanation. Further, previous studies have not reported any IBD within regions centred around introduction sites within the range (Rollins et al. 2009), indicating significant exchange between localities and suggesting that the historic introduction regime is an unlikely factor driving primary population substructure.

#### Comparison of Population structure of Australian and North American starlings

Replicated starling introductions in Australia and North America allow us to explore how invasive populations may develop diverse genetic characteristics in response to different environments. Although analyses indicate clearly defined environmental regions across the starling’s range in Australia, environmental parameters examined in this study explained little of the genome-wide genetic structure observed in Australian starling populations (Supplementary Material, Fig. S3). A non-significant environmental association using the genome-wide SNP dataset may indicate that high levels of vagility are obscuring signals of IBE. It is also possible that previously observed morphological association with environmental gradients is due to plasticity or is driven by genetic change at only a small number of loci. This later explanation is more likely, as geographic distance showed a strong relationship with, and explained a large amount of the variation in, genetic difference between collection localities, and also correlated with environmental distance. These findings contrast with similar analysis conducted on the North American starlings, which found geographic distance was a poor predictor of genetic differentiation across the species range, and that environment may be playing a greater role in population similarity and clustering (Hofmeister, et al. 2019). Some but not all starling populations migrate across North America, and this variation in migratory strategy may result in a stronger signature of IBE.

Collection localities within regions of Australia show low levels of genetic differentiation, nevertheless F_ST_ values are higher than those values reported in the larger (square km coverage) North American starling distribution (Hofmeister, et al. 2019). This further supports the notion that there are greater constraints to gene flow across the Australian continent than in North America. We note that the genetic division between VIC and NSW reported by both genome-wide and outlier loci SNP data sets straddles two key environmental barriers; the Great Dividing Range (an area of high elevation), and inland NSW (extreme aridity). Despite the fact that this is a highly dispersive species, it is possible that these environmental features restrict gene flow. While further work is required to examine the underlying causes for starling demographic and invasion history differences, this system demonstrates both the malleability of invasion regimes and the importance of studying replicated global invasions.

### Putatively adaptive variation

There is an expectation that local adaptation in highly mobile species (such as starlings) will be rare (Kvistad, et al. 2015). High levels of gene flow are often seen in highly mobile species, which acts to homogenise genetic differences between populations, while stabilising selection results in the loss of rare variants (North, et al. 2011). However, locally adapted variants have been identified in species that are highly mobile (Nielsen, et al. 2009; Limborg, et al. 2012; Bourret, et al. 2013; Hess, et al. 2013; Matala, et al. 2014; Milano, et al. 2014; Benestan, et al. 2015), and in introduced species (Rohfritsch, et al. 2013; Chown, et al. 2015), which indicates that local adaptation can occur despite high levels of gene flow.

Over all analyses, several hundred unique candidate loci (375) for local adaptation were identified, and from this, a total of 25 proteins were identified (Supplementary Material, Table S3). Support for the occurrence of adaptive selection across the starling’s range comes from the functional nature of the polymorphisms detected (Supplementary Material, Table S2) (Dudaniec, et al. 2018; Leydet, et al. 2018). Many of the biological functions are shared with the functions of putative loci under selection identified from the 15,038 SNPs found in in the North American starling study (Hofmeister, *et al*. 2019), and two genes were reported as candidates for selection in both studies (*COL18A*, tissue morphogenesis; *DEF6*, cell morphology regulation; Supplementary Material, Table S3). This overlap adds strength to the notion that both invasive populations are undergoing concurrent rapid adaptation.

Interestingly, we identify many more SNPs that are associated with temperature than precipitation or aridity. In contrast, Hofmeister *et al*. (2019) found a more even distribution across a similarly diverse array of environmental predictors in environmental association tests. While an excess of temperature associated SNPs may be (in part) a result of linkage, one plausible biological explanation may be that annual temperatures of the starling’s native range are more similar to that of North America than Australia. Further, as invasive starling populations are strongly associated with suburban and rural areas (Zufiaurre, et al. 2016), it is possible that temperature may exert a selective pressure that is less affected by human-mediated environmental disturbances than other selective forces (e.g. aridity). Further analysis on the role anthropogenic land alteration has on starling spread and evolution is an important future direction for this research.

Among the other proteins associated with Bio05 are a range of biological functions plausibly linked to temperature, for instance a calcium channel protein (CACNA1C; Supplementary Material, Table S2) that plays an important role in synaptic plasticity and hence behaviour, with modification of this gene being associated with changed learning strategies in mice (Koppe, et al. 2017). The importance of behavioural flexibility is a well-established in invasion biology (Wright, et al. 2010), and starlings are known for high relative brain sizes, a measure that has been linked to greater invasion success (Sol and Lefebvre 2000).

There was a particularly strong association between temperature and the candidate SNP associated with the genomic region that codes for Tcdd-Inducible Poly(Adp-Ribose) Polymerase (TIPARP). This protein is involved in pathways controlling androgen and estrogen metabolic process (and hence sexual characteristics), as well as the development of blood, skeleton and various organ structures (Supplementary Material, Table 3) (Ame, et al. 2004). Given the large number of functions associated with TIPARP, there could be multiple reasons it is associated with temperature, including variation in exposure to xenobiotic chemicals across a temperature gradient, or because temperature variation is highly dependent on latitude, and correlated with other seasonal fluctuations, both of which cue sexual maturation in starlings (Dawson 2005).

However, closer examination of the plots of allelic frequency across environmental variables reveals that environmental associations may be confounded by local signatures of founding populations. Sampling sites that are fixed for a single allele may not be randomly distributed, but rather may appear exclusively (or predominantly) within a single genetic group. For example, the locus strongly associated with Bio15 (Fig. 5c) is fixed in eight localities, six of which are in the northern Australia genetic cluster. Similar patterns can be seen in Fig. 5b, while Fig. 5a represents a site where fixation does not occur within the same subpopulation. It is possible that the alternative allele is not present in those areas due to founding effects at introduction, and therefore there may be other loci linked to environmental variables whose relationship is obscured by the legacy of limited diversity in some founding populations. While genomic approaches provide unprecedented sensitivity and power for testing for signatures of selection, caution should be applied when interpreting results for invasive species, or populations that have recently undergone rapid expansion, as neutral loci may then mimic signature of selection (Lotterhos and Whitlock 2014) (Riquet, et al. 2013). Characterising the native range genomic landscape for an invasive species would provide invaluable information to help discern the nature of polymorphisms within their invasive range, allowing us to identify if they are existing rare polymorphisms, or novel in the invasive range. Lastly, our study demonstrates the utility of using multiple approaches when searching for loci putatively under selection. We report some overlap between candidate loci identified through the three methods we used. However, many of the loci were reported for only one of the exploratory methods, and all of the annotated proteins identified were unique to their identification method.

In Australian starling populations, we showed that geographic distance plays a strong role in the genetic structure of populations, and that environmental barriers are likely reinforcing the observed patterns of genetic structure. Our genetic evidence also suggests that the contemporary Australian population of common starlings is most likely made up of two large and spatially distinct populations, plus one smaller southwestern NSW group, and one smaller genetic group existing at the Western-most range edge. These patterns are very different from those observed in a similar study on North American starlings, a demonstration that this species has the capacity for adaptive invasion strategies. Strong IBD and environmental associations for individual loci are highly suggestive of adaptive genetic variation, and doubtless these selective pressures will continue to drive population differentiation.

## Acknowledgements

PC was supported by an ARC Future Fellowship (FT0991420). LR was supported by an ARC Discovery Early Career Research Award (DE150101393). Thank you to Katherine L. Buchanan and Katie Hyma for their assistance with this project and manuscript. Thank you to Natalie R. Hofmeister for the useful and insightful discussions regarding these data. Thank you to the four anonymous reviewers for the constructive comments they provided, helping us to improve our manuscript. AC would like to acknowledge the lives of the 499 starlings who were killed and used for this research, and acknowledge an additional 236 starlings who were killed in conjunction with this project.

## Data Accessibility

Scripts and data are archived on GitHub, and will be made public upon publication: https://github.com/katarinastuart/Sv1_StarlingGBS

