## Supplementary Material for "Signatures of selection in a recent invasion reveals adaptive divergence in a highly vagile invasive species"

**Supplementary File**

### Filering parameters

Three variable datasets were created with a range of SNP minimum minor allele frequencies (mnMAF - 0.01, 0.05 and 0.10) to test the effects of different filtering parameters and identify a suitable dataset for downstream population genetic analysis. Sites that did not have a 'PASS' filter score were discarded, adapters were removed and all reads were trimmed to 64 bp.

All filtering from this point was done using the software VCFtools (Danecek et al. 2011). Blank and duplicated samples were removed from each dataset, the duplicated sample with the greatest number of reads was kept. Genotypes were filtered by depth where 1, 3 and 5 reads per individual were kept. Potentially overrepresented genotypes with removed from all datasets by filtering out genotypes with a maximum genotype depth of 60; approximately 10X the average genotype depth. All loci that were missing in more than 90% of samples were removed and subsequently all samples that had less than 10% of the total number of loci were removed. Each dataset was then filtered based on 9 levels of sample 'missingness', from 10% to 90% at 10% intervals. This filtering process included levels of mnMAF, genotype depth and sample 'missingness' and resulted in 81 differently filtered datasets (Supplementary Fig. S1).

The number of samples and loci retained across the 81 datasets after filtering ranged from 494 – 505 samples and 0 – 134,259 loci (see Supplementary Fig. S1). Each data set was tested for Hardy Weinberg Equilibrium (HWE) using VCFtools, and after false discovery rate correction of p-values, any SNPs out of HWE were removed. A fastSTRUCTURE analysis was performed on each of the 81 filtered data sets to determine the effect of different filtering options on estimates of genetic structure.

### Population structure analysis

*Population structure based on genome-wide SNP loci*

For the FASTSTRUCTURE analysis, a plot of $K_{\varepsilon}^{*}$ values from K = 1 to K = 10 was created (Fig. S2). To ensure that it was reasonable to use the genome-wide dataset without removing SNPs contributing to non-neutral variation, we created a neutral dataset that had all outlier loci removed (see 'detection and characterisation of adaptive variation' subsection below) and identified the number of K (Fig. S2b). Because we found no difference in the K plots between the genome-wide and neutral datasets, we used the former for downstream analysis. After identifying the appropriate K for the dataset, FASTSTRUCTURE was run 100 times with a logistic prior and the optimal K. The meanQ values from the 25 runs with the highest marginal likelihood values were averaged. Modified R scripts written by Mikhail Matz (https://goo.gl/NRmdCP) were used to identify and calculate meanQ values from the top 25 logistic runs. We created structure plots using average meanQ values (Fig. S3).

*Population structure based on outlier SNP loci*

A FASTSTRUCTURE analysis was run using only the outlier loci to identify if there were distinct genetic clusters associated with these putatively adaptive loci (Fig. S2c). To validate that the outlier dataset was better than random, a random subset of an equivalent number of SNPs was selected from the genome-wide dataset and a FASTSTRUCTURE analysis conducted (Fig. S2c).


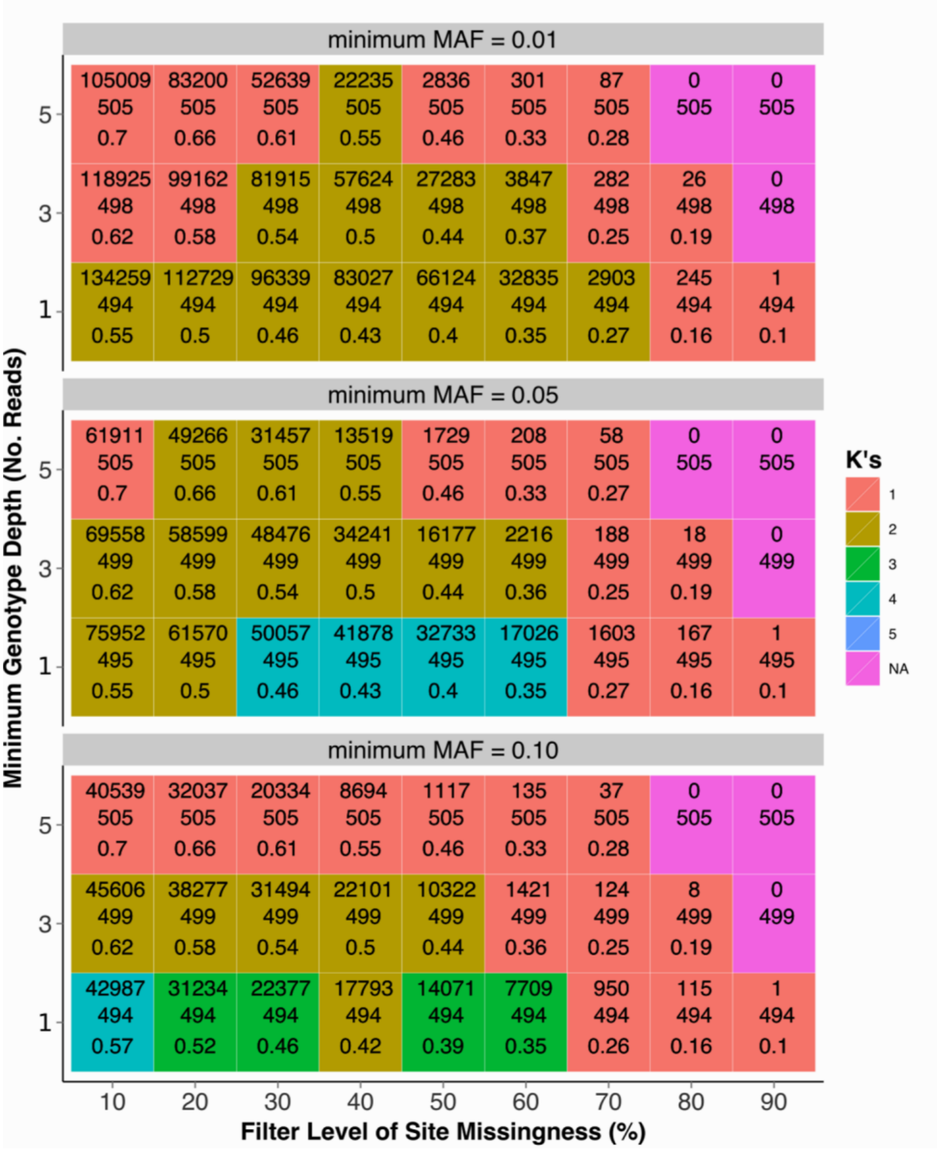


**Fig. S1.** Heat map showing population structure of Australian starlings using a range of 81 different combinations of filtering parameters including genotype depth, minimum minor allele frequency (MAF) and level of sample missingness allowed. Square colour indicates the number of K as identified by fastSTRUCTURE analysis.

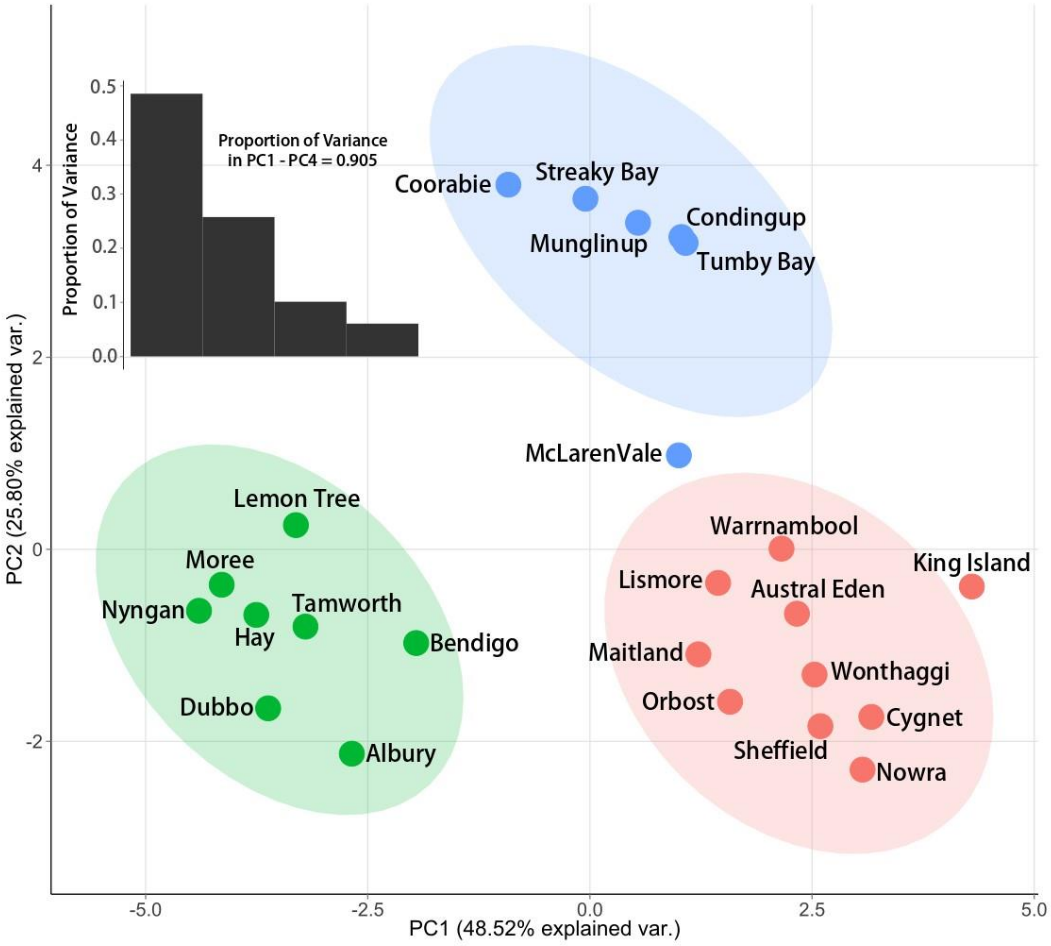


**Fig. S2.** PCA plot of Australian starling collection localities in environmental space. PC1 and PC2 were derived from 15 environmental variables. The PC1 axis describes variation in temperature and temperature seasonality and the PC2 axis describes variation in precipitation and the seasonality of precipitation. The colour of a point indicates the environmental cluster it falls within, and the coloured ellipse represents the data ellipse for each cluster as defined by the function stat_ellipse. Blue indicated arid localities, green indicated semi-arid localities, and blue indicates non-arid localities. A histogram of eigenvalues indicates the proportion of variance described by PC1 – PC4.

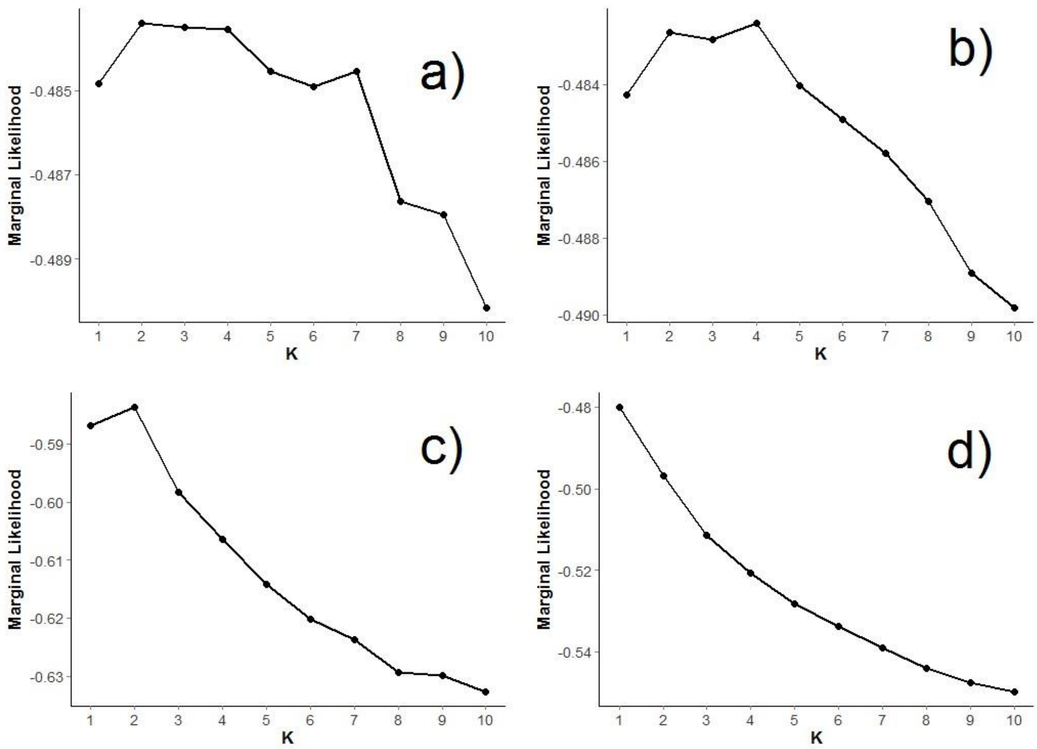


**Fig. S3.** Plot identifying the best value of K from multiple fastSTRUCTURE runs of Australian starling SNP data with K = 1 to 10. The K value with the highest marginal likelihood value identifies the number of clusters in the dataset that best describe genetic structure. Each panel describes the results for a different dataset: a) genome-wide dataset (N = 16,177 SNPs), b) neutral dataset with outlier loci removed (N = 16, 088), c) outlier dataset (N = 89 SNPs), and d) random subset of loci dataset (N = 89 SNPs).

**Table S1.** Statistics and PC loadings from a PCA analysis of 15 environmental variables taken from Australian starling collection sites.

| **Variable** | **PC1** | **PC2** | **PC3** | **PC4** |
|---|---|---|---|---|
| **Elevation** | -0.286 | -0.103 | 0.011 | 0.042 |
| **Distance to Introduction** | -0.034 | 0.325 | 0.075 | 0.705 |
| **Distance to Coast** | -0.327 | -0.152 | -0.058 | -0.014 |
| **Aridity** | 0.289 | -0.279 | 0.192 | -0.007 |
| **Mean NDVI** | 0.213 | -0.334 | 0.170 | 0.336 |
| **Variability in Day Length** | 0.221 | -0.161 | -0.575 | 0.109 |
| **Bio02** (Mean Diurnal Range) | -0.350 | 0.023 | 0.163 | 0.165 |
| **Bio03** (Isothermality) | 0.132 | 0.380 | 0.123 | 0.395 |
| **Bio04** (Temperature Seasonality) | -0.353 | -0.139 | 0.084 | -0.043 |
| **Bio05** (Max Temperature of Warmest Month) | -0.343 | 0.045 | 0.263 | -0.035 |
| **Bio06** (Min Temperature of Coldest Month) | 0.241 | 0.292 | 0.244 | -0.185 |
| **Bio07** (Temperature Annual Range) | -0.360 | -0.086 | 0.100 | 0.049 |
| **Bio13** (Precipitation of Wettest Month) | 0.203 | -0.202 | 0.558 | -0.092 |
| **Bio14** (Precipitation of Driest Month) | 0.122 | -0.427 | 0.230 | 0.206 |
| **Bio15** (Precipitation Seasonality) | 0.087 | 0.405 | 0.207 | -0.319 |
| **Standard Deviation** | 2.698 | 1.965 | 1.231 | 0.953 |
| **Proportion of Variance** | 0.485 | 0.258 | 0.101 | 0.06061 |
| **Cumulative Proportion** | 0.485 | 0.743 | 0.844 | 0.905 |

**Table S2.** $F_{ST}$ values for all pairwise comparisons between collection localities of Australian starlings. Values below the diagonal are pairwise $F_{ST}$ values calculated from the genome-wide dataset (N = 16,177 SNPs), whereas values above the line were calculated using the outlier dataset (N = 89 SNPs). The heat map colour scale from clear to dark blue indicates low to high $F_{ST}$ values for each data set respectively. Collection localities are arranged from west to northeast following geographic distribution. Red numbers identify those pairwise $F_{ST}$ values that were considered non-significant ($p > 0.05$) after false discovery rate correction for multiple comparisons. Column labels correspond to collection localities, but have been shortened so that all values fit on one page.

| Collection Localities | Mun. | Con. | Coo. | SB | TB | McL. | War. | Ben. | Won. | Orb. | KI | Shef. | Cyg. | Alb. | Hay | Dub. | Nyn. | Now. | Mait. | Aust. | Tam. | Mor. | Lis. | Lem. |
|---|---|---|---|---|---|---|---|---|---|---|---|---|---|---|---|---|---|---|---|---|---|---|---|---|
| **Munglinup** | - | 0.274 | 0.269 | 0.320 | 0.272 | 0.255 | 0.268 | 0.224 | 0.261 | 0.255 | 0.221 | 0.258 | 0.264 | 0.211 | 0.206 | 0.248 | 0.253 | 0.282 | 0.295 | 0.253 | 0.305 | 0.231 | 0.284 | 0.216 |
| **Condingup** | 0.090 | - | 0.094 | 0.151 | 0.123 | 0.107 | 0.075 | 0.079 | 0.116 | 0.091 | 0.045 | 0.100 | 0.079 | 0.112 | 0.087 | 0.182 | 0.159 | 0.202 | 0.240 | 0.173 | 0.231 | 0.165 | 0.155 | 0.150 |
| **Coorabie** | 0.079 | 0.029 | - | 0.174 | 0.034 | 0.038 | 0.121 | 0.083 | 0.146 | 0.141 | 0.075 | 0.099 | 0.069 | 0.106 | 0.088 | 0.142 | 0.147 | 0.183 | 0.220 | 0.167 | 0.218 | 0.122 | 0.140 | 0.100 |
| **StreakyBay** | 0.113 | 0.044 | 0.022 | - | 0.189 | 0.128 | 0.227 | 0.159 | 0.231 | 0.192 | 0.158 | 0.166 | 0.159 | 0.161 | 0.153 | 0.185 | 0.209 | 0.239 | 0.216 | 0.164 | 0.247 | 0.191 | 0.174 | 0.149 |
| **TumbyBay** | 0.076 | 0.030 | 0.004 | 0.017 | - | 0.007 | 0.152 | 0.127 | 0.196 | 0.194 | 0.096 | 0.100 | 0.069 | 0.134 | 0.122 | 0.196 | 0.196 | 0.238 | 0.254 | 0.216 | 0.273 | 0.169 | 0.207 | 0.169 |
| **McLarenvale** | 0.081 | 0.033 | 0.003 | 0.026 | 0.002 | - | 0.115 | 0.102 | 0.176 | 0.159 | 0.072 | 0.068 | 0.063 | 0.116 | 0.109 | 0.171 | 0.164 | 0.202 | 0.221 | 0.172 | 0.227 | 0.150 | 0.179 | 0.132 |
| **Warnambool** | 0.073 | 0.034 | 0.018 | 0.034 | 0.020 | 0.020 | - | 0.056 | 0.060 | 0.083 | 0.017 | 0.110 | 0.091 | 0.057 | 0.054 | 0.134 | 0.129 | 0.153 | 0.201 | 0.149 | 0.166 | 0.136 | 0.121 | 0.118 |
| **Bendigo** | 0.075 | 0.037 | 0.019 | 0.050 | 0.022 | 0.023 | 0.005 | - | 0.030 | 0.070 | 0.024 | 0.073 | 0.063 | 0.045 | 0.022 | 0.082 | 0.091 | 0.123 | 0.152 | 0.085 | 0.133 | 0.085 | 0.091 | 0.049 |
| **Wonthaggi** | 0.078 | 0.041 | 0.026 | 0.042 | 0.026 | 0.030 | 0.007 | 0.007 | - | 0.051 | 0.060 | 0.152 | 0.122 | 0.048 | 0.055 | 0.120 | 0.125 | 0.138 | 0.176 | 0.107 | 0.167 | 0.120 | 0.124 | 0.083 |
| **Orbost** | 0.080 | 0.042 | 0.020 | 0.048 | 0.023 | 0.025 | 0.011 | 0.011 | 0.012 | - | 0.062 | 0.131 | 0.112 | 0.048 | 0.058 | 0.124 | 0.097 | 0.135 | 0.165 | 0.107 | 0.133 | 0.126 | 0.106 | 0.092 |
| **KingIsland** | 0.076 | 0.034 | 0.015 | 0.030 | 0.018 | 0.018 | 0.006 | 0.009 | 0.010 | 0.012 | - | 0.046 | 0.065 | 0.036 | 0.026 | 0.092 | 0.088 | 0.123 | 0.155 | 0.085 | 0.125 | 0.100 | 0.073 | 0.062 |
| **Sheffield** | 0.089 | 0.046 | 0.026 | 0.040 | 0.027 | 0.025 | 0.026 | 0.028 | 0.028 | 0.028 | 0.022 | - | 0.051 | 0.105 | 0.061 | 0.120 | 0.130 | 0.164 | 0.157 | 0.125 | 0.177 | 0.122 | 0.115 | 0.091 |
| **Cygnet** | 0.088 | 0.044 | 0.022 | 0.042 | 0.026 | 0.024 | 0.024 | 0.029 | 0.030 | 0.027 | 0.023 | 0.004 | - | 0.087 | 0.060 | 0.126 | 0.128 | 0.187 | 0.206 | 0.153 | 0.205 | 0.128 | 0.121 | 0.098 |
| **Albury** | 0.077 | 0.044 | 0.020 | 0.078 | 0.024 | 0.023 | 0.011 | 0.009 | 0.017 | 0.011 | 0.014 | 0.029 | 0.030 | - | 0.017 | 0.055 | 0.052 | 0.068 | 0.102 | 0.059 | 0.081 | 0.046 | 0.071 | 0.045 |
| **Hay** | 0.077 | 0.039 | 0.019 | 0.060 | 0.023 | 0.022 | 0.008 | 0.007 | 0.013 | 0.011 | 0.013 | 0.026 | 0.027 | 0.002 | - | 0.036 | 0.050 | 0.066 | 0.087 | 0.042 | 0.084 | 0.032 | 0.049 | 0.038 |
| **Dubbo** | 0.086 | 0.047 | 0.025 | 0.073 | 0.027 | 0.029 | 0.020 | 0.019 | 0.026 | 0.019 | 0.023 | 0.031 | 0.032 | 0.005 | 0.005 | - | 0.026 | 0.041 | 0.040 | 0.025 | 0.024 | 0.025 | 0.030 | 0.022 |
| **Nyngan** | 0.080 | 0.038 | 0.022 | 0.039 | 0.020 | 0.025 | 0.017 | 0.016 | 0.018 | 0.015 | 0.019 | 0.022 | 0.025 | 0.012 | 0.009 | 0.005 | - | 0.039 | 0.054 | 0.028 | 0.040 | 0.035 | 0.032 | 0.037 |
| **Nowra** | 0.091 | 0.049 | 0.030 | 0.057 | 0.031 | 0.034 | 0.028 | 0.026 | 0.032 | 0.026 | 0.028 | 0.036 | 0.035 | 0.018 | 0.016 | 0.007 | 0.007 | - | 0.053 | 0.049 | 0.048 | 0.041 | 0.058 | 0.058 |
| **Maitland** | 0.099 | 0.055 | 0.035 | 0.066 | 0.036 | 0.038 | 0.032 | 0.031 | 0.038 | 0.029 | 0.032 | 0.039 | 0.039 | 0.021 | 0.021 | 0.008 | 0.009 | 0.004 | - | 0.036 | 0.043 | 0.070 | 0.058 | 0.051 |
| **Austral** | 0.081 | 0.041 | 0.021 | 0.048 | 0.023 | 0.026 | 0.016 | 0.014 | 0.021 | 0.016 | 0.018 | 0.027 | 0.027 | 0.011 | 0.011 | 0.005 | 0.003 | 0.006 | 0.006 | - | 0.053 | 0.039 | 0.049 | 0.034 |
| **Tamworth** | 0.088 | 0.043 | 0.026 | 0.038 | 0.026 | 0.031 | 0.021 | 0.022 | 0.024 | 0.019 | 0.021 | 0.027 | 0.026 | 0.021 | 0.018 | 0.008 | 0.001 | 0.004 | 0.006 | 0.001 | - | 0.048 | 0.037 | 0.043 |
| **Moree** | 0.084 | 0.044 | 0.022 | 0.060 | 0.025 | 0.028 | 0.020 | 0.017 | 0.025 | 0.018 | 0.020 | 0.030 | 0.031 | 0.011 | 0.010 | 0.003 | 0.003 | 0.003 | 0.004 | 0.002 | 0.002 | - | 0.068 | 0.025 |
| **Lismore** | 0.081 | 0.039 | 0.022 | 0.043 | 0.024 | 0.026 | 0.014 | 0.014 | 0.018 | 0.014 | 0.015 | 0.027 | 0.027 | 0.011 | 0.009 | 0.007 | 0.004 | 0.007 | 0.008 | 0.001 | 0.002 | 0.002 | - | 0.049 |
| **Lemon** | 0.078 | 0.040 | 0.021 | 0.047 | 0.023 | 0.025 | 0.017 | 0.016 | 0.016 | 0.016 | 0.018 | 0.024 | 0.027 | 0.010 | 0.010 | 0.005 | 0.001 | 0.007 | 0.008 | 0.004 | 0.003 | 0.002 | 0.003 | - |

**Table S3**. List of proteins that were matched to outlier loci, as calculated by Bayescan, BayEnv2, and RDA analyses of Australian starling SNPs. Method describes which analysis was used to identify each locus. Variable identifies the environmental variable or RDA axis with which each locus was associated for the environmental association approached (BayEnv2 and RDA). Values associated with BayEnv2 identified proteins indicates the magnitude of the association, while the loading associated with RDA identified proteins indicated both the magnitude and direction (positive or negative) of the association. Gene provides the Uniprot ID, and name provides the submitted name for each protein. Biological function provides an abbreviated list of the main biological functions associated with each protein, as reported on Uniprot.

| Method | Variable | Value/Loading | Gene | Name | Biological Function |
|---|---|---|---|---|---|
| Bayescan | | | OBSCN | Cytoskeletal Calmodulin And Titin-Interacting Rhogef | G protein-coupled receptor signaling pathway, positive regulation of apoptotic process, sarcomere organization |
| | | | ADGRD2 | Adhesion G Protein-Coupled Receptor D2 | Adenylate cyclase-activating G protein-coupled receptor signaling pathway, cell surface ad G protein-coupled receptor signaling pathways |
| | - | - | LOC106855771 | C-Factor-Like | Uncharacterized protein |
| | | | C1QC | Complement Component 1, Q Subcomponent, C Chain | Complement cascade (microbe response), immune response, negative regulation of macrophage and granulocyte |
| | | | LOC106858259 | Alpha-2-Macroglobulin-Like Protein 1 | Regulation of endopeptidase activity |
| | | | CHD2 | Chromodomain Helicase Dna Binding Protein 2 | Chromatin organization, muscle organ development, regulation of transcription by RNA polymerase II |
| | | | LOC106860795 | Zonadhesin-Like | Cell adhesion |
| BayEnv2 | Aridity | 4.14 | HSPA9 | Heat Shock Protein Family A | Cellular response to heat and unfolded proteins, protein refolding, and negative regulation of apoptotic process |
| | Bio05 | 3.8 | STAB1 | Stabilin 1 | Cell-cell signaling and adhesion, bacterium and inflammation response |
| | | 3.43 | CACNA1C | Calcium Channel, Voltage-Dependent, L Type, Alpha 1C Subunit | Calcium ion transmembrane transport, and cardiac conduction |
| | | 5.13 | CEP63 | Centrosomal Protein 63Kda | Cell devision (specifically G2/M transition of mitotic cell cycle), centriole replication, and spindle assembly |
| | | 6.6 | ATAD2B | Atpase Family, Aaa Domain Containing 2B | Negative regulation of chromatin silencing, and positive regulation of transcription by RNA polymerase II |

| | | | | | |
|---|---|---|---|---|---|
| | | 18.1 | TIPARP | Tcdd-Inducible Poly(Adp-Ribose) Polymerase (Tiparp) | Androgen and estrogen metabolic process, blood, skeleton and various organ structure formation/development |
| | | 3.63 | CORIN | Corin, Serine Peptidase | Female pregancy, peptide hormone processing, and blood pressure regulation |
| | | 6.4 | TMED6 | Transmembrane P24 Trafficking Protein 6 | Golgi organization, endoplasmic reticulum to Golgi vesicle-mediated transport, intracellular protein transport |
| | | 3.47 | COL18A1 | Collagen, Type Xviii, Alpha 1 | Angiogenesis, animal organ morphogenesis, positive regulation of endothelial cell apoptotic process |
| | | 4.42 | ATP8B1 | Atpase, Aminophospholipid Transporter, Class I, Type 8B, Member 1 | Ion transmembrane transport, Golgi organization, negative regulation of transcription |
| | Bio15 | 3.21 | CTRL | Chymotrypsin-Like | Protein catabolic process |
| RDA | Axis 2 | 0.070958 | TRAF1 | Tnf Receptor Associated Factor 1 | Apoptotic process, protein-containing complex assembly, protein-containing complex assembly |
| | Axis 2 | -0.06662 | LOC106856597 | Short Transient Receptor Potential Channel 2-Like | Acrosome reaction, calcium ion transmembrane transport, mating behavior, sex discrimination |
| | Axis 2 | 0.066728 | P3H3 | Prolyl 3-Hydroxylase 3 | Collagen biosynthetic and metabolic process, negative regulation of cell population proliferation |
| | Axis 3 | -0.06581 | DEF6 | Guanine Nucleotide Exchange Factor | Regulate cell morphology, T helper cells development and/or activation |
| | Axis 3 | -0.06815 | LOC106851981 | Inositol 1,4,5-Trisphosphate Receptor-Interacting Protein-Like 1 | Encodes protein that enhances the sensitivity of ITPR to intracellular calcium signaling |
| | Axis 3 | -0.06945 | HINFP | Histone H4 Transcription Factor | DNA repair, establishment of protein localization, in utero embryonic development, negative regulation of gene expression and transcription |
| | Axis 3 | -0.06577 | HCN3 | Hyperpolarization Activated Cyclic Nucleotide Gated Potassium Channel 3 | Cellular response to dopamine, regulation of ion transmembrane transport, response to cisplatin |
| | Axis 3 | -0.06604 | SNX21 | Sorting Nexin Family Member 21 | Protein transport |

**Function for scraping google for elevation data.**

This function scrapes googles elevation API for elevation data at lon, lat coordinates that you feed it. This function was working in 2014 but may no longer work based on changes to googles API.

```
googEl <- function(locs)  {
	require(RJSONIO)
	locstring <- paste(do.call(paste, list(locs[, 2], locs[, 1], sep=',')), collapse='|')
	u <- sprintf('http://maps.googleapis.com/maps/api/elevation/json?locations=%s&sensor=false', locstring)
	res <- fromJSON(u)
	out <- t(sapply(res[[1]], function(x) {
		c(x[['location']]['lat'], x[['location']]['lng'], x['elevation'], x['resolution'])
	}))
	rownames(out) <- rownames(locs)
	return(out)
}
```

**Function for imputing genotype for RDA.**

```
#calculate probabilities of the three genotypes for each column (SNP)
n <- nrow(SNPdata)
p_0 <- apply(SNPdata, 2, function(x){sum(x == 0, na.rm = T)/(n - sum(is.na(x)))})
p_1 <- apply(SNPdata, 2, function(x){sum(x == 1, na.rm = T)/(n - sum(is.na(x)))})
p_2 <- apply(SNPdata, 2, function(x){sum(x == 2, na.rm = T)/(n - sum(is.na(x)))})
p <- data.frame(p_0, p_1, p_2)

#make a table for indices of missing genotypes
indices <- which(is.na(SNPdata), arr.ind = T)

#replace missing genotypes by sampling from (0,1,2) based on probabilities given in table p
for (i in nrow(indices)) {
```

```
  x <- NA_indices[i, ]
  SNPdata [x[1], x[2]] <- sample(c(0:2), 1, replace = T, prob = p[x[2], ])
}
```
